# Site-1 Protease is a negative regulator of sarcolipin promoter activity

**DOI:** 10.1101/2025.02.25.639963

**Authors:** Isha Sharma, Meredith O. Kelly, Katelyn Hanners, Ella S. Shin, Muhammad G. Mousa, Shelby Ek, Gretchen A. Meyer, Rita T. Brookheart

**Affiliations:** John T. Milliken Department of Medicine, Division of Nutritional Science and Obesity Medicine, Washington University School of Medicine, St. Louis, MO, 61110, USA; Program in Physical Therapy, Washington University School of Medicine, St. Louis, MO 63110, USA; Departments of Orthopaedic Surgery and Neurology, Washington University School of Medicine, St. Louis, MO 63110, USA

**Keywords:** ATF6, S1P, calcium, calcineurin, sarcolipin, endoplasmic reticulum, CREB

## Abstract

The timed contraction and relaxation of myofibers in tissues such as the heart and skeletal muscle occurs via the tightly regulated movement of calcium ions into and out of the sarcoplasmic reticulum (SR). In skeletal muscle, this phenomenon enables humans to exercise, perform day-to-day tasks, and to breathe. Sarcolipin, a small regulatory protein, prevents calcium ions from entering the SR by binding to and inhibiting SERCA, contributing to myofiber contraction. Disruptions in sarcolipin expression are implicated in the pathophysiology of obesity and musculoskeletal disease. However, the mechanisms regulating sarcolipin expression are not clearly understood. We recently showed that Site-1 Protease (S1P) is a regulator of skeletal muscle function and mass. Here, we report that deleting S1P in mouse skeletal muscle increases sarcolipin expression, without impacting calcium SR flux. In cultured cells, S1P negatively regulates sarcolipin by activating the transcription factor ATF6, which inhibits basal- and calcineurin-stimulated sarcolipin promoter activity. We identified a cAMP response element binding protein (CREB) binding site on the sarcolipin promoter that is necessary for promoter activation, and show that in muscle, CREB binds to the sarcolipin promoter and that this binding is enhanced when S1P is deleted. These discoveries expand our knowledge of S1P biology and the mechanisms controlling calcium regulatory genes.

## INTRODUCTION

In skeletal muscle and the heart, the transient movement of calcium ions from sarcoplasmic reticulum (SR) calcium stores to the cytosol results in myofiber contraction and, to relax these fibers, the active transport of calcium ions back into the SR^1^. This transport of calcium ions relies heavily on the sarco/endoplasmic reticulum Ca2+ ATPase pump (SERCA)^1,2^. SERCA activity is inhibited by the direct binding of two small proteins, sarcolipin and phospholamban^3^. Sarcolipin is predominantly expressed in skeletal muscle, while phospolamban is highly expressed in cardiac muscle^3^. Both proteins inhibit SERCA, which leads to calcium accumulation in the cytosol and the relaxation of myofibers – underscoring their importance for proper muscle mechanics, and in the context of skeletal muscle, organismal movement^4–6^. While phospolamban is implicated in cardiac myopathies, disrupted sarcolipin expression has predominately been linked to the pathophysiology of Duchenne muscular dystrophy (DMD), obesity, and type 2 diabetes^7–13^. Despite the importance of sarcolipin to maintain normal skeletal muscle function, movement, and its link to disease, little is understood about how sarcolipin is regulated^14,15^.

Site-1 Protease (S1P; also known as subtilisin/kexin-isozyme 1 or PCSK8) is as Golgi-resident protease that regulates cellular homeostasis by cleaving and activating a variety of transcription factors (REF). S1P has been classically studied in liver and bone as a regulator of cellular lipid homeostasis and the unfolded protein response^16–19^. We recently demonstrated that S1P controls skeletal muscle function and size and that excessive S1P activity impairs exercise tolerance in a person with an S1P gain-of-function mutation, emphasizing the importance of S1P in mammalian muscle biology^20,21^. Here, we show that skeletal muscle-specific S1P knockout mice (S1P^smKO^) have increased sarcolipin expression in multiple muscle groups yet have normal skeletal muscle SR calcium flux and contractile function. Our data indicate that S1P inhibits sarcolipin expression by preventing the binding of CREB to the sarcolipin promoter and that activation of the S1P-substrate ATF6 acts as an inhibitor of calcineurin-dependent sarcolipin transcription. Our study unveils a function for S1P in the control of sarcolipin and identifies a mechanism by which this occurs.

## RESULTS

### Sarcolipin expression is increased in S1P depleted skeletal muscle

We recently reported that S1P is a key component of skeletal muscle biology and identified 75 significantly differentially expressed genes in the gastrocnemius of S1P skeletal muscle-specific knockout (S1P^smKO^) mice using bulk RNA sequencing^20,21^. Our RNASeq data showed that sarcolipin transcript levels were significantly increased in the gastrocnemius of S1P^smKO^ mice compared to WT mice (Figure 1A). We validated these data by measuring sarcolipin mRNA and protein levels by qPCR and western blotting, respectively and confirmed that sarcolipin expression is increased in the gastrocnemius of S1P^smKO^ mice compared to WT mice (Figure 1B-C). Muscle group-specific differences in sarcolipin expression have been reported and we previously showed that S1P can alter expression of genes in a muscle group-specific manner, particularly in muscle that contains glycolytic fibers like the gastrocnemius and quadraceps^20,22^. Thus, we measured the levels of sarcolipin protein in other muscle groups and observed increased sarcolipin levels in the quadriceps, but not in the soleus, a muscle composed primarily of oxidative fibers (Figure 1D and Supplemental Figure 1A). Together these data indicate that S1P negatively regulates sarcolipin expression in a subset of skeletal muscle groups that contain glycolytic fibers such as gastrocnemius and quadriceps.

**Figure 1.**
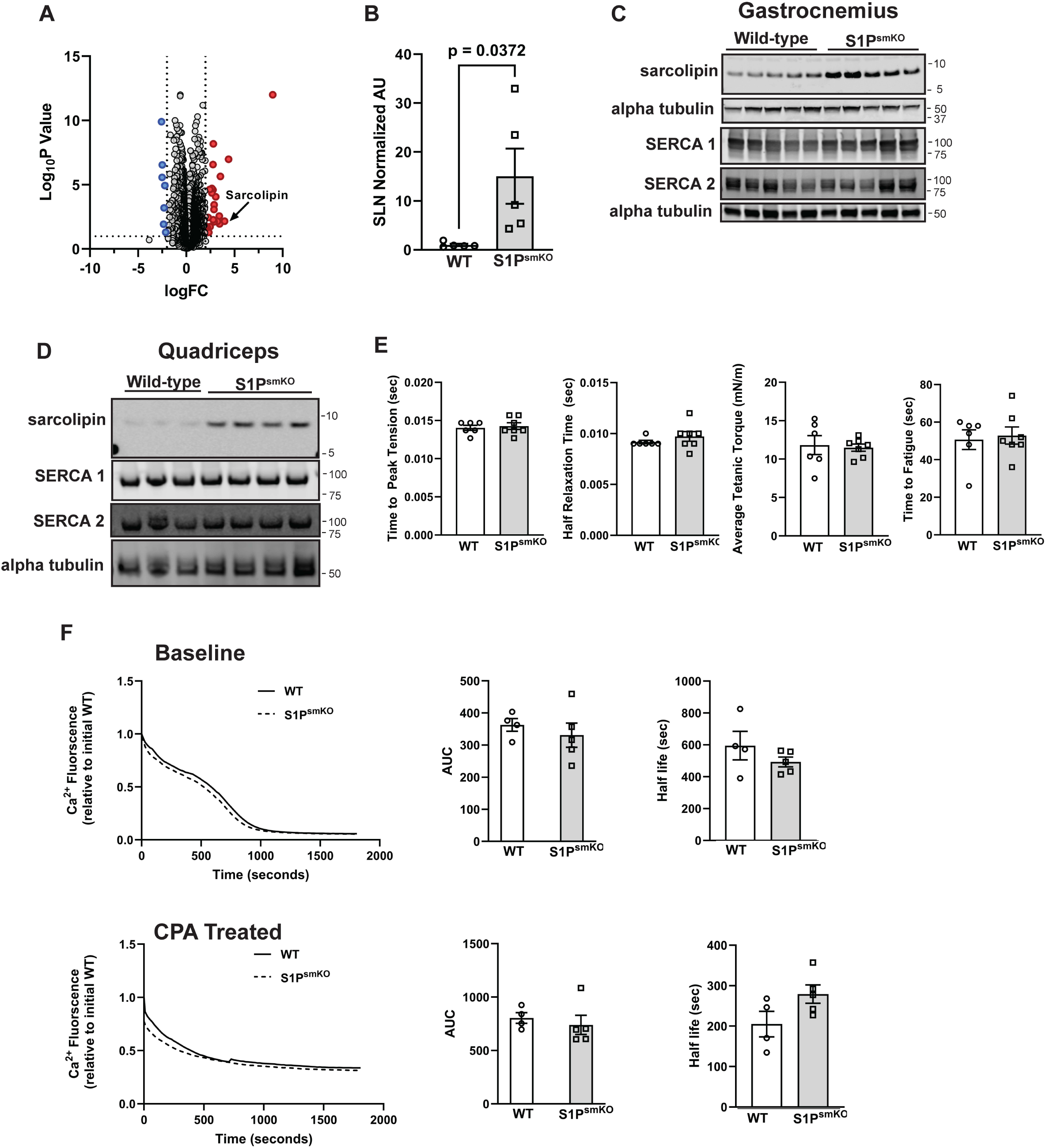
Deletion of S1P in skeletal muscle increases sarcolipin transcription and blunts SR calcium flux. (A) Volcano plot of genes identified from RNA-Seq as significantly differentially increased (red dots) and decreased (blue dots) in gastrocnemius of S1P^smKO^ mice relative to WT mice. Sarcolipin is indicated. n=4 per genotype. (B) qPCR of sarcolipin mRNA expression in gastrocnemius of S1P^smKO^ and WT mice. n=5 per group. Western blot of sarcolipin and SERCA protein levels in (C) gastrocnemius and (D) quadriceps of S1P^smKO^ and WT mice. n = 4-5 per group. (E) Peak tension of twitch and tetanic contractions and (F) time to fatigue of S1P^smKO^ and WT gastrocnemius. Rates of ATP-stimulated free calcium SR uptake with area under the curve (left) and uptake half-life (right) at baseline (F) and with CPA treatment (G). Data are reported as ± SEM.

### S1P deletion in skeletal muscle does not impact contractile function or SR calcium flux

Disruptions in skeletal muscle calcium regulators, such as sarcolipin, compromise contractile function in muscles like the soleus and plantaris ^23–26^. Because the gastrocnemius and quadricep muscles of S1P^smKO^ mice have high levels of sarcolipin compared to WT mice, we examined whether this increase in sarcolipin correlated with changes in muscle contractile function. To do this, we measured *in vivo* plantarflexor torque of S1P^smKO^ and WT mice. The mouse plantar flexor muscles consist of gastrocnemius, soleus, and plantaris muscles, of which the gastrocnemius accounts for 70.6 % of total muscle mass^27^. Twitch kinetics (peak twitch torque, time to peak torque and half-relaxation time) were unchanged between KO and WT muscles. Additionally, we observed no differences in isometric tetanic contraction or fatigability between KO and WT mice. Together these data indicate that the measured change in sarcolipin expression in the gastrocnemius of S1P^smKO^ mice did not manifest in differences peak torque or fatiguability (Figure 1E).

Sarcolipin inhibits SERCA activity in skeletal and cardiac muscle and overexpressing sarcolipin in these tissues has been shown to decrease rates of SR calcium uptake^4,5,28^. Based on these data and the increased sarcolipin levels in S1P^smKO^ gastrocnemius and quadricep muscles, we examined whether SR calcium uptake was altered in S1P^smKO^ gastrocnemius. To test this, we quantified SR ATP-stimulated free calcium uptake in gastrocnemius homogenates of S1P^smKO^ and WT mice and observed no difference in the rates of SR calcium uptake or half-life between KO and WT tissues. To specifically examine SERCA-dependent ATP-stimulated free calcium uptake, we measured uptake in homogenates treated with the SERCA-specific inhibitor cyclopiazonic acid (CPA) or vehicle and noted that rates of uptake and half-life were unchanged between KO and WT gastrocnemius muscles (Figure 1F). Together these data indicate that in the gastrocnemius of S1P^smKO^ mice, an increase in sarcolipin protein is not associated with changes in SR calcium flux or contractile function.

### S1P inhibits sarcolipin promoter activity

S1P controls an array of transcription factors that drive the expression of genes required for cellular homeostasis^29^. However, it has been reported that some of these transcription factors also inhibit gene expression by suppressing promoter activity^30,31^. Because sarcolipin expression is increased in S1P^smKO^ muscles, we examined whether S1P was indirectly inhibiting sarcolipin promoter activity. To measure sarcolipin transcription, we created a sarcolipin luciferase reporter construct that contains 1,339 base pairs of the mouse sarcolipin promoter upstream of a cDNA encoding a luciferase reporter (Figure 2A). We validated this construct in murine C2C12 cells by co-expressing it with constitutively active calcineurin (CnA), a known activator of the sarcolipin promoter^14^ and observed a 51-fold increase in sarcolipin promoter activity compared to empty vector control cells (Figure 2B-C). To further validate the sarcolipin reporter construct, we tested the ability of the calcineurin inhibitor RCAN1 to suppress sarcolipin promoter activity. Indeed, overexpressing RCAN1 completely inhibited calcineurin-dependent sarcolipin promoter activity, consistent with previous studies (Figure 2B-C)^14^.

**Figure 2.**
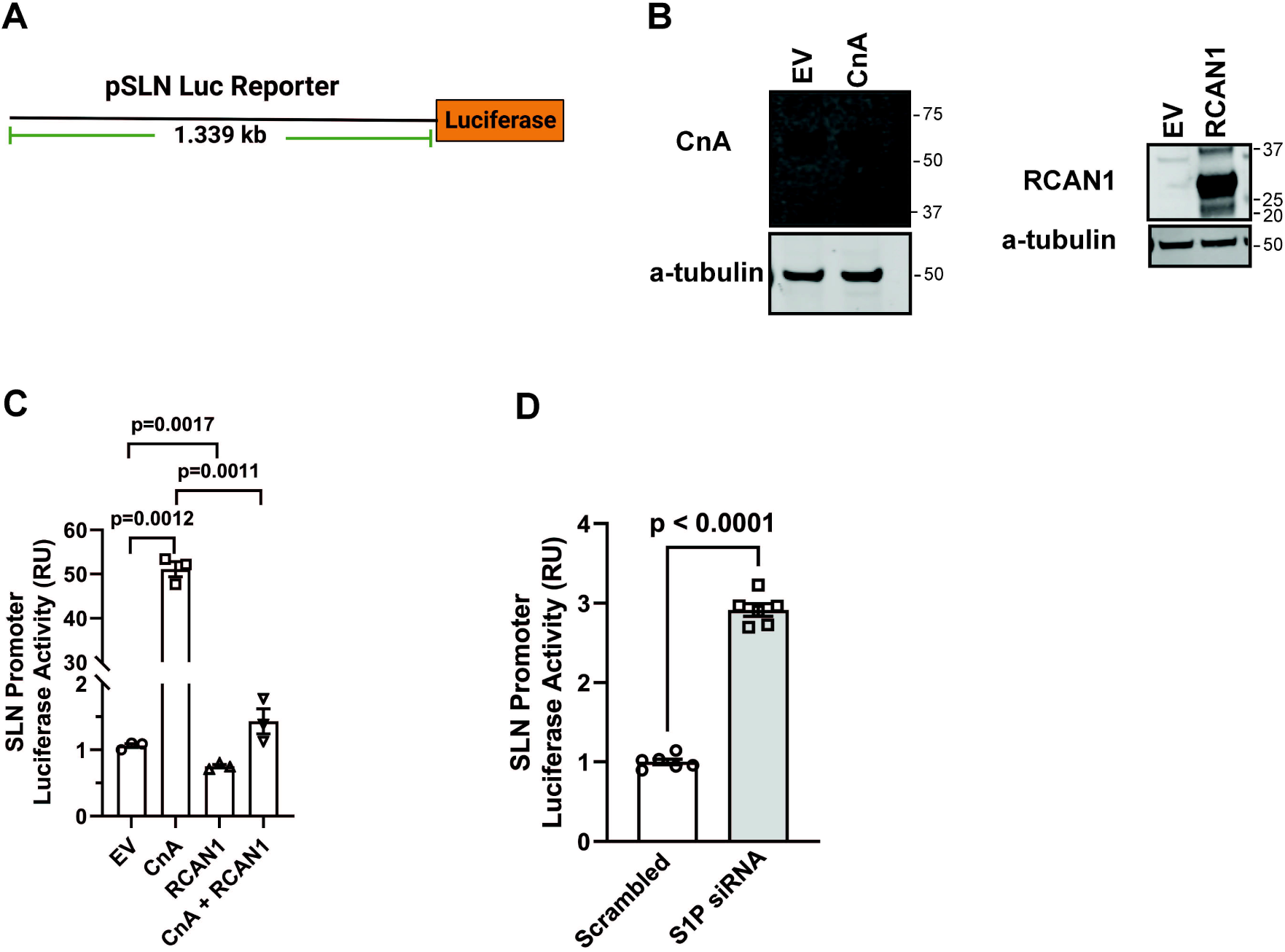
S1P inhibits basal and calcineurin-dependent sarcolipin transcription. (A) Schematic of sarcolipin luciferase reporter construct containing 1.339 kb of the mouse sarcolipin promoter. (B) Western blot of calcineurin (left) and RCAN1 (right) protein levels in C2C12 cells transiently transfected with EV, CnA, or RCAN1 plasmids. Representative blots of n=3. (C) Quantification of sarcolipin promoter activity in C2C12 cells transfected with the indicated plasmids. n=5 per group. (D) Sarcolipin promoter activity in scrambled and S1P knockdown C2C12 cells. n=5-6 per group. (E) Western blot of EV and S1P-KDEL protein levels in cells transfected with the indicated plasmids. Representative blots of n=3. (F) Sarcolipin promoter activity in cells expressing EV, S1P-KDEL, and CnA. n=5 per group. EV, empty vector; CnA, constitutively active calcineurin. Data are reported as ± SEM.

With this validated sarcolipin promoter reporter system, we next examined sarcolipin promoter activity in the absence or presence of S1P. We transiently knocked down S1P in C2C12 cells using small interfering RNA (siRNA) oligos, as we previously reported^20^. Cells transfected with a scrambled siRNA served as a negative control. Scrambled and S1P knockdown cells were transfected with the sarcolipin promoter construct, and promoter activity was measured. Relative to negative control cells, depletion of S1P in C2C12 cells increased sarcolipin promoter activity over 2-fold (Figure 2D). These data indicate that loss of S1P de-represses sarcolipin promoter activity.

### ATF6 restores repression of the sarcolipin promoter in S1P depleted cells

S1P is known to activate the transcription factor ATF6, which can selectively activate or repress promoter activity.^30–33^ Unlike the other transcription factors that S1P activates, ATF6 inhibits calcineurin, a protein that drives sarcolipin promoter activity (Figure 2C)^14,34,35^. To determine if the S1P-substrate ATF6 is an inhibitor of the sarcolipin promoter, C2C12 cells were co-transfected with the sarcolipin promoter construct and an expression vector to express a truncated form of ATF6 (nATF6) or empty vector control. Since nATF6 lacks the transmembrane domain that tethers ATF6 to the ER and Golgi, nATF6 constitutively localizes to the nucleus and controls gene expression in an unregulated manner^36^. In the presence of nATF6, sarcolipin promoter activity was inhibited compared to cells transfected with an empty vector control (Figure 3B). Additionally, when cells were depleted of S1P, nATF6 still inhibited sarcolipin promoter activity (Figure 3C). These data suggest that the S1P-substrate ATF6 is an inhibitor of sarcolipin promoter activation.

**Figure 3.**
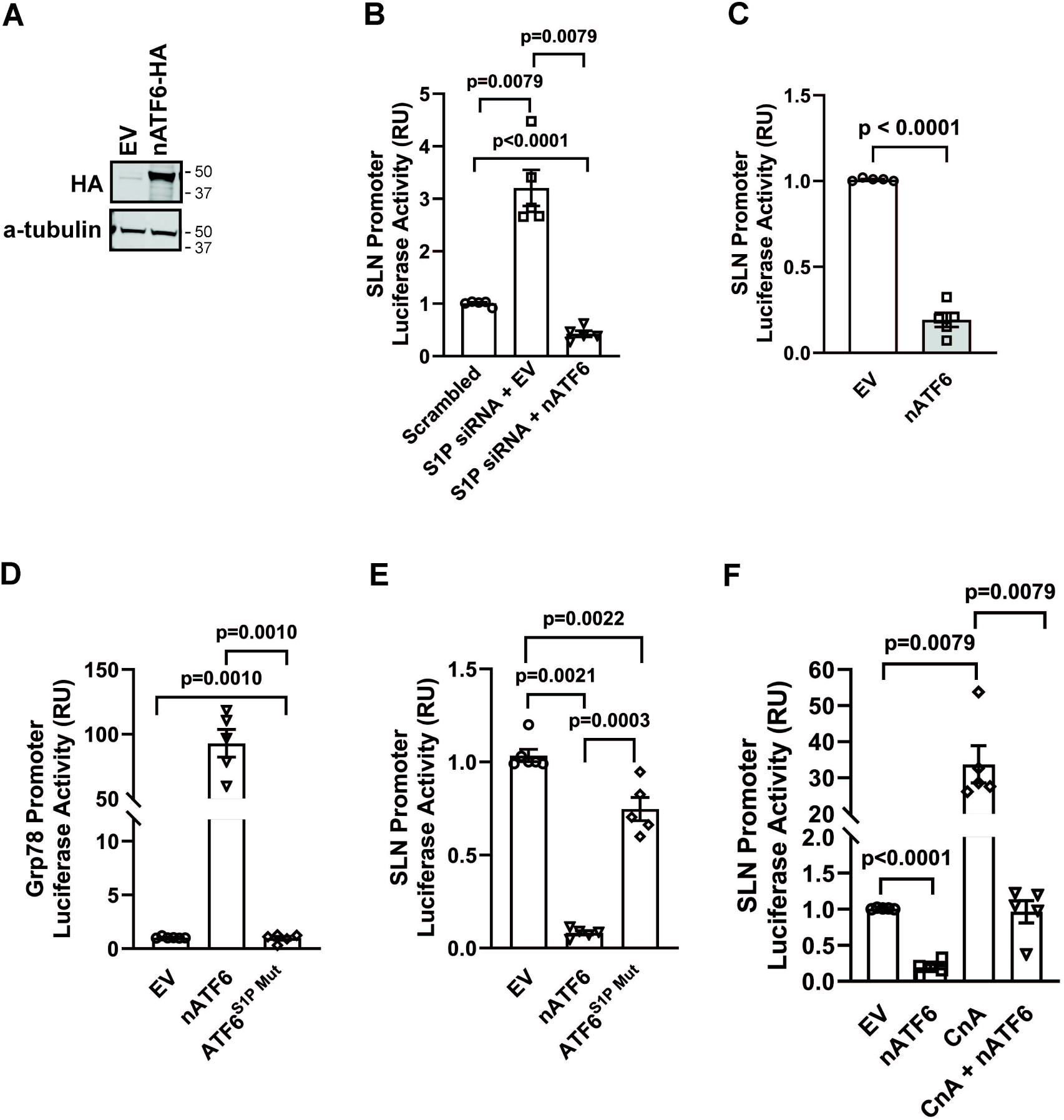
S1P inhibits sarcolipin transcription by activating ATF6. (A) Western blot of cells transfected with EV or constitutively active ATF6 (nATF6-HA). Representative blot of n=3. (B) ATF6-dependent transcription reporter activity in cells expressing EV or nATF6. n=5 per group. (C) Sarcolipin promoter activity in scrambled +EV, S1P knockdown + EV, and S1P knockdown + nATF6 cells. n=5 per group. (D) Western blot of ATF6^S1Pmut^ protein levels in cells transfected with EV or ATF6^S1Pmut^ plasmids. Representative blot of n=3. (E) ATF6-dependent transcription reporter activity in cells expressing EV or ATF6^S1Pmut^. n=5 per group. Sarcolipin promoter activity in cells expressing (F) EV or ATF6^S1Pmut^ and cells expressing (G) EV, CnA, and nATF6 as indicated. n=5 per group. EV, empty vector; CnA, constitutively active calcineurin. Data are reported as ± SEM.

S1P cleaves ATF6, triggering the subsequent translocation of ATF6 to the nucleus where it can control promoter activity^34,36^. However, reports have shown that ATF6 can also be activated independently of S1P cleavage, and that this leads to selective but still significant changes in gene transcription^34^. To determine if ATF6’s inhibition of the sarcolipin promoter was dependent on S1P cleavage, we measured sarcolipin promoter activity in C2C12 cells expressing either empty vector, nATF6 (constitutively active), or an ATF6 mutant lacking the S1P cleavage site (ATF6^S1Pmut^)^36^. While nATF6 completely suppressed sarcolipin promoter activity, ATF6^S1Pmut^ caused a slight, but significant reduction in sarcolipin promoter activity compared to control cells (Figure 3E). As a control, we tested the ability of ATF6^S1Pmut^ to activate an ATF6-responsive luciferase reporter construct, which it did not (Figure 3D). These data indicate that the S1P-dependent cleavage of ATF6 is a necessary precursor for ATF6’s inhibition of the sarcolipin promoter.

### ATF6 suppresses calcineurin-dependent sarcolipin promoter activity

Since ATF6 inhibits, while calcineurin activates, the sarcolipin promoter (Figure 2C and 3B)^14^ and ATF6 is known to suppress calcineurin-dependent signaling^34,35^, we hypothesized that the reciprocal control of the sarcolipin promoter by ATF6 and calcineurin were interconnected. To test this, we examined whether ATF6 prevented calcineurin from activating the sarcolipin promoter. Overexpressing constitutively active calcineurin (CnA) increased sarcolipin promoter activity (Figure 3F)^14^; however, co-expressing active ATF6 (nATF6) with CnA decreased calcineurin-dependent sarcolipin promoter activation down to basal levels (Figure 3F). These data indicate that nATF6 completely blocks calcineurin-stimulated activation of the sarcolipin promoter, placing ATF6 downstream of calcineurin signaling in this context.

In the heart, ATF6 inhibits calcineurin activation, possibly by increasing expression of the calcineurin inhibitor RCAN1^35^. We and others have shown that RCAN1 blocks calcineurin-stimulated sarcolipin promoter activity (Figure 2C)^14^. This would suggest that S1P controls sarcolipin expression via an ATF6-RCAN1 axis that targets calcineurin. To test this possibility, we used two different siRNAs to deplete RCAN1 in C2C12 cells, and measured sarcolipin promoter activity in the presence of nATF6 (Supplemental Figure 1B-C). Despite depleting RCAN1 with two different siRNAs, active ATF6 still inhibited calcineurin-stimulated sarcolipin promoter activity (Supplemental Figure 1C), suggesting that ATF6 disrupts calcineurin signaling independently of RCAN1.

### Sarcolipin promoter activity is driven by CREB in vitro and in vivo

To determine which portion of the sarcolipin promoter is important for basal and calcineurin-stimulated promoter activity, we generated a series of truncation mutants of our sarcolipin luciferase reporter construct and examined their transcriptional activity at baseline and in response to constitutively active calcineurin (Figure 4A-B). At baseline, all truncation mutants had reduced promoter activity compared to the full-length promoter (-1339) (Figure 4B). In the presence of active calcineurin, sarcolipin promoter activity increased for all mutant promoters relative to their untreated controls, with mutant -19 exhibiting the lowest increase (Figure 4B). These data suggest that nucleotides -989 through -19 of the sarcolipin promoter are essential for promoter activation both at baseline and in response to calcineurin stimulation.

**Figure 4.**
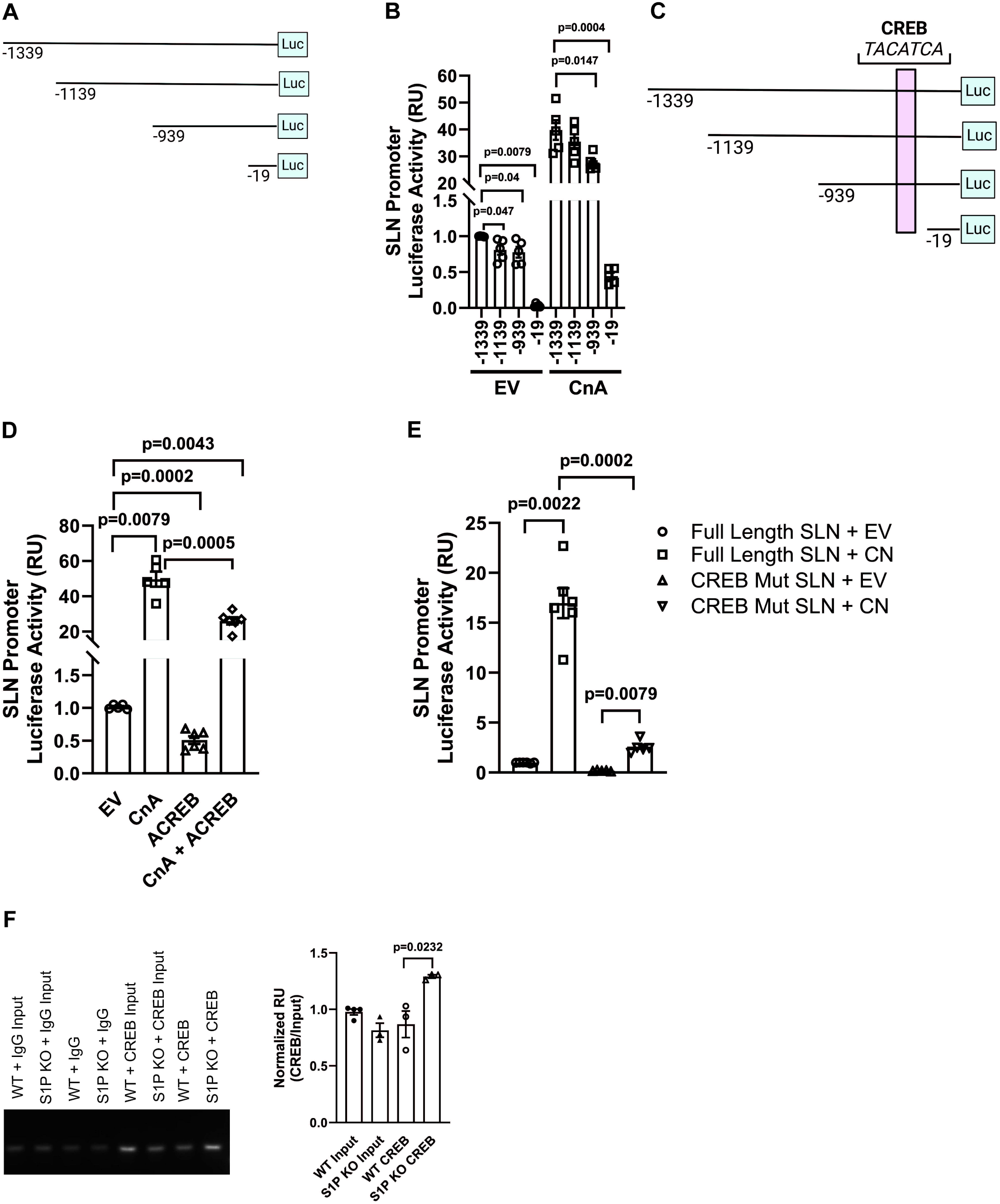
CREB binds to the sarcolipin promoter and is necessary to activate calcineurin-dependent sarcolipin transcription. (A) Schematic of sarcolipin promoter reporter construct deletions used in the study. Figures denote the number of base pairs upstream of the transcriptional start site. (B) Sarcolipin promoter activity in cells expressing the indicated truncated versions of the sarcolipin promoter reporter construct. Cells were transfected with EV or CnA. (C) Diagram of predicted CREB DNA binding site on the mouse sarcolipin promoter. (D) Sarcolipin promoter activity in cells transfected with EV, CnA, or CREBA as indicated. (E) Sarcolipin promoter activity in cells transfected with SLN FL or SLN CREB mut and with either EV or CnA, as indicated. (F) Chromatin immunoprecipitation of gastrocnemius of WT and S1P^smKO^ mice quantified (graph, n=4) and representative agarose gel of ChIP qPCR products (n=3). SLN FL, full length sarcolipin promoter construct; SLN CREB mut, sarcolipin promoter construct with CREB binding site mutated; EV, empty vector; CnCA, constitutively active calcineurin, CREBA, dominant negative CREB. Data are reported as ± SEM.

Rotter *et al*. identified two predicted CREB binding sites on the sarcolipin promoter, one of which is positioned between nucleotides -989 and -19 (Figure 4C)^14^. We hypothesized that CREB is an important component of sarcolipin promoter activity and may control the promoter at this predicted CREB site. To test this hypothesis, we first examined whether a dominant negative CREB (ACREB)^37^ can inhibit sarcolipin promoter activation at baseline and in the presence of active calcineurin. At baseline, expression of ACREB in C2C12 cells decreased sarcolipin promoter activity by 50 % compared to cells expressing empty vector (Figure 4D). In the presence of active calcineurin, ACREB inhibited sarcolipin promoter activity by 48 % compared to cells expressing active calcineurin alone (Figure 4D). These data indicate that a functional CREB protein is a necessary factor for activating the sarcolipin promoter both at baseline and in response to calcineurin.

We next examined whether the CREB binding site between -989 and -19 identified by Rotter *et al.* is required for sarcolipin promoter activiation^14^. We mutated this CREB binding site in our full-length sarcolipin luciferase reporter construct and examined promoter activity at baseline and in the presence of active calcineurin (Figure 4C). Mutation of the CREB binding site inhibited basal promoter activity by 87% relative to control cells expressing an intact full-length promoter construct (Figure 4E). In the presence of active calcineurin, mutation of the CREB binding site inhibited promoter activity by 85% compared to calcineurin-treated cells expressing the intact full-length promoter (Figure 4E). These data suggest that the CREB binding site on the sarcolipin promoter is required for promoter activity both at baseline and in response to calcineurin. These cell culture studies show that CREB and its promoter binding site are required for sarcolipin promoter activity. We extended these studies *in vivo* and examined the binding of CREB to the endogenous sarcolipin promoter of S1P^smKO^ and WT gastrocnemius. Compared to WT gastrocnemius, muscle from S1P^smKO^ mice had increased binding of CREB to the sarcolipin promoter (Figure 4F). Together our *in vitro* and *in vivo* data identify sarcolipin as a CREB target gene in skeletal muscle and that calcineurin is a driver of CREB-dependent sarcolipin promoter activation.

## DISCUSSION

In the present study, we define several novel and complex regulatory circuits controlling the expression of sarcolipin in skeletal myocytes. We show that S1P negatively regulates sarcolipin expression in skeletal muscle and define the mechanisms at play. Interestingly, we report that S1P’s influence on sarcolipin transcription does not impact contractile function or SERCA-dependent SR calcium flux. This latter finding suggests that sarcolipin activity is controlled by mechanisms that are beyond transcriptional regulation. Indeed, post-translational modifications are a known regulatory mechanism for controlling sarcolipin and SERCA^38–40^. One group reported that phosphorylation of sarcolipin by the kinase STK16 inhibits its ability to suppress SERCA^40^. Thus, it is possible that the increased levels of sarcolipin protein observed in S1P^smKO^ skeletal muscles remains inactive due to the presence of an inhibitory protein modification – a possibility that our lab is exploring.

Several studies have reported that increased sarcolipin levels in the soleus muscle correlate with defective SR calcium uptake and contractile dysfunction ^23–26^. Interestingly, reports using a diverse range of mouse models have shown that sarcolipin levels are increased in the soleus (a muscle composed primarily of oxidative muscle fibers), but not in mixed muscle groups like the gastrocnemius, which has a combination of oxidative and glycolytic muscle fibers^41,42^. S1P^smKO^ mice have increased sarcolipin in the gastrocnemius and quadriceps, another mixed fiber-type muscle group (Figure 1C and 1D). Yet, we observed no increase in sarcolipin expression in the soleus of S1P^smKO^ mice (Supplemental Figure 1A). It is well documented that the type of muscle group dictates the function, gene expression, cellular calcium flux, and metabolism of the muscle ^23,43–47^. Indeed, our recent publication showed that S1P controls muscle gene expression and size in a muscle-type-dependent manner^20^. Our data coupled with those from previous groups suggests that the regulation of sarcolipin expression and its influence on SR calcium flux is muscle-type dependent^41,42^. Our ongoing work is focused on teasing apart the mechanisms by which gene expression and muscle function are differentially regulated based on muscle type.

In the heart, ATF6 suppresses calcineurin signaling through the protein RCAN1^35^. Our data suggests that ATF6 inhibits calcineurin-stimulated sarcolipin promoter activity independently of RCAN1. Our approach used siRNAs to deplete cells of RCAN1, it is possible that the remaining amounts of RCAN1 or other RCAN isoforms were sufficient for ATF6 to block calcineurin signaling and prevent sarcolipin promoter activity. It is also possible that ATF6 inhibits sarcolipin promoter activity through an RCAN1-independent mechanism that involves CREB since ATF6 also functions as an inhibitor of CREB DNA-binding^48^.

In conclusion, our study uncovered a complex series of regulatory players that control sarcolipin expression in skeletal muscle. We show that S1P inhibits sarcolipin promoter activity through its activation of the transcription factor ATF6 and that ATF6 suppresses the sarcolipin promoter by blocking calcineurin-dependent promoter activation. We also identified a functional CREB-binding site on the sarcolipin promoter and showed that CREB binds to and is an activator of this promoter. This work contributes to our understanding of the dual function that transcription factors can play as both activators and suppressors of gene promoters. Sarcolipin is implicated in the pathogenesis of Duchenne muscular dystrophy (DMD) and metabolic disease, where it is suggested to play conflicting roles — high levels of sarcolipin are associated with negative DMD disease outcome, while high levels of sarcolipin correlates with improved muscle function in obese mice, but not in humans with obesity ^7–12^. The nature of this discrepancy is not clear, but understanding how sarcolipin expression and activity are controlled in skeletal muscle will provide the foundational knowledge upon which we begin to understand this gene’s importance in human disease.

## METHODS

### Animal Studies

All mouse studies were approved by the Institutional Animal Care and Use Committee of Washington University in St. Louis. S1P floxed and S1P skeletal muscle-specific knockout mice in the C57BL/6J background were previously described^16,20^. Littermates not expressing Cre recombinase (S1P floxed mice) were used as controls for all experiments. Mice were genotyped for the presence of Cre recombinase and floxed S1P allele using gene-specific primers, as described previously^20^. Studies were performed using male mice aged 12-14 weeks of age. Mice were fed a standard laboratory chow diet and group housed on a 12 h light/dark cycle.

### Plantarflexor Torque Testing

Contractile function of the plantarflexor muscle group (which consists of 70.6 % gastrocnemius muscle by mass) was assessed via stimulation of the tibial nerve as previously described^27,49^. Briefly, hindlimbs of anesthetized mice were shaved and a small ∼2mm long skin incision was made lateral to the knee where the sciatic nerve bifurcates into the peroneal and tibial branches. The peroneal branch was visualized by resecting the overlying musculature and then severed to avoid recruitment of the ankle dorsiflexors during stimulation. The skin was then closed with glue (Vetbond) and the mouse was transferred to a physiology testing rig (1300A Aurora Scientific), where it was positioned supine on a heated platform with the knee clamped and the ankle secured to a footplate attached to a dual mode ergometer. Two platinum needle electrodes were then inserted subcutaneously on either side of the tibial nerve. Optimal stimulation current was determined using a current sweep and then a twitch contraction was acquired to assess twitch kinetics (peak twitch torque, time to peak torque and half-relaxation time). Peak tetanic torque was determined as the average of three contractions at 150 Hz with 2 min rest times between. Following these measurements, a fatiguing protocol was delivered with 150 Hz tetanic contractions every 2 sec for 2 min. Time to fatigue was calculated as the time until peak torque dropped to 50% of the initial value.

### Constructs

pSLN-Luciferase (pSLN-Luc) construct was generated by GENEWIZ in which the first 1390 base pairs upstream of the mouse sarcolipin start site were cloned into pGL3-Basic (Promega) using NheI and BglIII restriction enzyme sites. Truncated and CREB mutant pSLN-Luc constructs were generated using QuikChange Lightening Site-Directed Mutagensis Kit (Agilent, Cat# 210518) as per manufacturer’s’ instructions and custom designed primers (Supplemental Table 1). Custom primers were designed with the Agilent QuikChange Primer Design Program (https://www.agilent.com/store/primerDesignProgram.jsp). Additional constructs used in this study are pCGN-ATF6 (1-373) (a gift from Ron Prywes - Addgene plasmid # 27173^50^), pEGFP-ATF6-(S1P-) (a gift from Ron Prywes - Addgene plasmid # 32956^36^); RCAN1 (a gift from Jeffery Molkentin to Addgene plasmid # 65413^51^); CnA^52^ ; ACREB^37^; S1P-KDEL (a gift from Peter Espenshade^53^); and pMAXGFP (Lonza).

### Cell culture

C2C12 cells were grown at 37 °C with 5% CO^2^ in DMEM supplemented with 10 % fetal bovine serum and 1% penicillin-streptomycin. For plasmid expression studies, cells were plated in 6-well plates at 1.2 x 10^5^ cells per well. After 24 h, cells were transfected with the indicated constructs using Lipofectamine 2000 (Life Technologies) as per manufacturer instructions. For siRNA studies, cells were transfected with RNAiMax (Life Technologies) as per manufacturer’s instructions. After 4 days post-transfection, cells were harvested using RIPA buffer (Sigma) for protein isolation or RNA-STAT 60 (Tel-TEST Inc) for RNA isolation.

### Luciferase assays

Cells were plated in 12-well plates at 5 x 10^4^ cells per well. After 24 h, cells were transfected with pSLN-Luc (either full length or truncated), pMAX-GFP, and additional constructs indicated in the figures using Lipofectamine 3000 (Life Technologies) as per manufacturer’s instructions. For studies using both constructs and siRNAs, cells were transfected 24 h after plating with the indicated constructs and siRNAs using Lipofectamine 2000 (Life Technologies) following manufacturer’s instructions for simultaneous transfection of cells with DNA and siRNAs. Four days post-transfection, luciferase assays were performed using One-Glo Luciferase Assay System (Promega, Cat # E6120) as per manufacturer’s instructions. Luminescence followed by GFP fluorescence were quantified with a BioTek plate reader.

### Immunoblotting

Skeletal muscle protein lysates were generated by homogenizing tissues in lysis buffer (20 mM Tris, 15 mM NaCl, 1 mM EDTA, 0.2% NP-40, and 10% glycerol) containing 2x Protease Complete cocktail tablet (Roche) and 1x Phosphatase Inhibitors (Roche) with 5 mm stainless steel beads in a TissueLyzer II (Qiagen). Protein lysates were rotated for 45 min at 4°C, centrifuged at 15,000 x g for 15 min at 4°C. For cultured cells, protein lysates were generated using RIPA buffer supplemented with 2x Protease Complete cocktail tablet (Roche) and 1X Phosphatase Inhibitors (Roche). Proteins were quantified by bicinchoninic acid assay (BCA, Pierce Biotechnology). Equal amounts of protein were resolved on a 4–12% BIS-Tris gradient gel (Invitrogen) with 1x MES running buffer and transferred to 0.2 µm nitrocellulose membrane (BioRad). Blots were probed with appropriate primary and secondary antibodies and proteins visualized by LI-COR Odyssey imaging system. The following antibodies were used in our studies at a 1:1000 dilution: alpha-tubulin (T5168, Sigma); SERCA1 (ab2819, Abcam); SERCA2 (ab3625, Abcam); sarcolipin (ABT13, Sigma); His-tag (2365S, Cell Signaling Technology); calcineurin (2614S, Cell Signaling Technology).

### Real Time Quantitative PCR

Total RNA was isolated from C2C12 cells and skeletal muscle with RNA STAT-60 (Tel-Test Inc) as per manufacturer’s instructions. For tissues, RNA was isolated by disrupting tissue in RNA STAT-60 using 5 mm steel beads (Qiagen) and a TissueLyser II (Qiagen). RNA was reverse transcribed into cDNA using the High-Capacity cDNA Reverse Transcription Kit (Applied Biosystems). Quantitative real-time PCR was performed using Power SYBR green (Applied Biosystems) and transcripts quantified on an ABI QuantiStudio 3 sequence detection system (Applied Biosystems). Data was normalized to 36B4 expression and results analyzed using the 2−ΔΔCt method and reported as relative units to controls. Primer sequences are listed in Table S1.

### Free Calcium Flux Studies

SERCA-mediated calcium uptake assay was performed as described^23,54^. Briefly, snap frozen mouse gastrocnemius muscles were cryopulverized, then homogenized with a cold steel bead in a Qiagen Tissuelyser II for 3 minutes at 30 Hz in 10 μL of Homogenizing buffer (250 mM sucrose, 5 mM HEPES, 0.2 mM PMSF, and 0.2% NaN3, pH 7.5) for each mg of sample. Samples were aliquoted and frozen to avoid more than 2 freeze-thaw cycles. Buffers were always kept on ice and samples were kept on ice until the initial read.

Samples were incubated with 1 μL of 2mM Fluo-4 (diluted in molecular grade water; ThermoFisher #F14200) in Ca2+ uptake buffer (200 mM KCl, 20 mM HEPES, 10 mM NaN3, 5 μM TPEN, 15 mM MgCl2, pH 7.0) for 10 minutes before reading on BioTek plate reader at 485nm excitation and 528nm emission. After the initial read, 4 μL 250 mM ATP (pH 7.0) was simultaneously added to each well using a multichannel pipettor to initiate the kinetic reaction and avoid row-dependent variance. Maximum fluorescence values were acquired by repeatedly adding 80 μL of 100 mM CaCl2 and reading until values stayed consistent. Minimum fluorescent values were acquired by repeatedly adding 30 μL 50mM EGTA (pH 7.0) to samples then reading until reaching plateau. Cyclopiazonic acid (2.2 μL10 mM, Sigma Aldrich #C1530) or equivalent volume of vehicle (DMSO) was added to each sample followed by addition of Fluo-4 and ATP and read as above to specifically determine SERCA-mediated calcium uptake.

Total sample protein amounts were quantified by Micro BCA Protein Assay Kit (ThermoFisher #23235). All initial and kinetic readers were normalized to total sample protein concentration, then to the initial read value (at t=0 seconds).

### Chromatin Immunoprecipitation (ChIP)

Snap-frozen mouse gastrocnemius tissue was pulverized and fixed with 1% formaldehyde solution containing phosphatase and protease inhibitors and rotated for 20 min at room temperature. Crosslinking was quenched with 2.5 M glycine solution with rotation for 5 min at room temperature. Samples were pelleted and washed twice in 1X PBS containing protease and phosphatase inhibitors. Samples were resuspended in 0.5% Igepal and 0.25% Triton X-100 lysis buffer with protease and phosphatase inhibitors followed by Dounce homogenization. Samples were pelleted and washed in Tris-HCl lysis buffer with protease and phosphatase inhibitors, spun down, and the resulting pellet was resuspended in low salt lysis buffer containing protease and phosphatase inhibitors followed by sonication to shear chromatin DNA. Optimal DNA shearing was confirmed by 1% agarose gel electrophoresis (250 bp – 1 kb smears). After adding 10% Triton X-100 to the sonicated lysate, samples were centrifuged to pellet cellular debris, and 50 µL of the supernatants were collected to serve as total genomic inputs. Samples were incubated with appropriate antibodies (1-10 µg of antibody per 25 ug of DNA) – normal IgG Rb (2729S, Cell Signaling Technology) or CREB antibody (9197S, Cell Signaling Technology) for 1 h at 4°C with rotation. Protein A/G magnetic beads (80105G, Invitrogen) were washed three times in 0.5% BSA blocking solution and resuspended in blocking solution with salmon sperm DNA (Sigma) with rotation for 30 min at room temperature. Beads were added to chromatin-antibody sample complexes and incubated overnight at 4°C with rotation. Samples were washed seven times with RIPA wash buffer containing protease and phosphatase inhibitors. Protein-DNA complexes were eluted from beads by incubating at 65°C for 15 min in Tris-HCl Elution buffer and further incubated at 65°C overnight to reverse protein-DNA crosslinks along with total genomic inputs. Samples were incubated with RNase A and then Proteinase K supplemented with 300 mM CaCl2 solution. DNA was isolated using the Qiagen Dneasy Blood and Tissue Kit (69594, Qiagen) as per manufacturer’s instructions. DNA was quantified by NanoDrop and qPCR was carried out as described above. Efficiency of sarcolipin DNA primers were confirmed by running a qPCR primer efficiency curve. Primer sequences are listed in Table S1.

*RNA-Sequencing Studies* of mouse gastrocnemius were performed as reported previously^20^. Accession number for the RNA-Seq data presented in this study was previously deposited at NCBI GEO under accession number GSE199014 located at https://www.ncbi.nlm.nih.gov/geo/query/acc.cgi?acc=GSE199014.

### Statistics

Normality of data distribution was examined with graphical inspection and the Shapiro-Wilk Test. Normally distributed data were analyzed by unpaired t-test and Welch’s Correction for unequal variances was applied when the homogeneity of variance assumption was violated. For data with unequal distribution, groups were compared using a Mann-Whitney Test. Data with small sample sizes (groups containing ≤ 4 observations) were analyzed with the Wilcoxon Test. Analysis was performed with GraphPad Prism Version 10.2.3 for Windows (Dotmatics). A p value ≤ 0.05 was considered statistically significant. Date are reported as ± SEM. Statistical details and the number of samples used in each study are indicated in the figure legends and figures.

### Data Availability

The RNA-seq datasets analyzed during the current study were originally reported in Mousa *et al.*^20^ and are deposited at NCBI GEO under accession number GSE199014 located at https://www.ncbi.nlm.nih.gov/geo/query/acc.cgi?acc=GSE199014.

## ACKNOWLEDGEMENTS

We thank Drs. Jay Horton at UT Southwestern for the S1P-floxed mouse strain. We also thank the Genome Technology Access Center (GTAC) at Washington University School of Medicine for help with RNASeq analysis. GTAC is partially supported by NCI Cancer Center Support grant P30 CA918342 to the Siteman Cancer Center and by ICTS/CTSA grant UL1TR002345 from the NCRR, a component of the NIH, and NIH Roadmap for Medical Research. Research reported in this publication was supported by a gift from Mrs. Carol MacCorkle, the Nutrition Obesity Research Center (NIH grant P30 DK056341), Washington University Musculoskeletal Research Center (NIH P30 AR074992), NIH grants K01 HL145326 and R25 DK132966. The content in this paper is solely the responsibility of the authors and does not necessary represent the official views of the NCRR or NIH.

## AUTHOR CONTRIBUTIONS

Conceptualization, R.T.B.; methodology, R.T.B. and G.A.M.; formal analysis, R.T.B., I.S., K.H., E.S.S., and M.O.K.; investigation, R.T.B., I.S., K.H., E.S.S., and M.O.K.; writing – original, R.T.B.; writing – review & editing, R.T.B. and G.A.M.; funding acquisition, R.T.B.; supervision, R.T.B. and G.M.

## REFERENCES

1. MacLennan, D.H. (2000). Ca2+ signalling and muscle disease. Eur. J. Biochem. 267, 5291–5297. 10.1046/J.1432-1327.2000.01566.X.

2. Periasamy, M., and Kalyanasundaram, A. (2007). INVITED REVIEW SERCA PUMP ISOFORMS: THEIR ROLE IN CALCIUM TRANSPORT AND DISEASE. Muscle Nerve 35, 430–442. 10.1002/mus.20745.

3. Shaikh, S.A., Sahoo, S.K., and Periasamy, M. (2016). Phospholamban and sarcolipin: Are they functionally redundant or distinct regulators of the Sarco(Endo)Plasmic Reticulum Calcium ATPase? J. Mol. Cell. Cardiol. 91, 81–91. 10.1016/j.yjmcc.2015.12.030.

4. Fajardo, V.A., Rietze, B.A., Chambers, P.J., Bellissimo, C., Bombardier, E., Quadrilatero, J., and Tupling, A.R. (2017). Effects of sarcolipin deletion on skeletal muscle adaptive responses to functional overload and unload. https://doi.org/10.1152/ajpcell.00291.2016 313, C154–C161. 10.1152/AJPCELL.00291.2016.

5. Babu, G.J., Zheng, Z., Natarajan, P., Wheeler, D., Janssen, P.M., and Periasamy, M. (2005). Overexpression of sarcolipin decreases myocyte contractility and calcium transient. Cardiovasc. Res. 65, 177–186. 10.1016/j.cardiores.2004.08.012.

6. Fajardo, V.A., Bombardier, E., McMillan, E., Tran, K., Wadsworth, B.J., Gamu, D., Hopf, A., Vigna, C., Smith, I.C., Bellissimo, C., et al. (2015). Phospholamban overexpression in mice causes a centronuclear myopathy-like phenotype. Dis. Model. Mech. 8, 999–1009. 10.1242/dmm.020859.

7. Paran, C.W., Verkerke, A.R.P., Heden, T.D., Park, S., Zou, K., Lawson, H.A., Song, H., Turk, J., Houmard, J.A., and Funai, K. (2015). Reduced efficiency of sarcolipin-dependent respiration in myocytes from humans with severe obesity. Obesity 23, 1440–1449. 10.1002/oby.21123.

8. Balakrishnan, R., Mareedu, S., and Babu, G.J. (2022). Reducing sarcolipin expression improves muscle metabolism in mdx mice. 10.1152/ajpcell.00125.2021.

9. Voit, A., Patel, V., Pachon, R., Shah, V., Bakhutma, M., Kohlbrenner, E., Mcardle, J.J., Dell’italia, L.J., Mendell, J.R., Xie, L.-H., et al. Reducing sarcolipin expression mitigates Duchenne muscular dystrophy and associated cardiomyopathy in mice. 10.1038/s41467-017-01146-7.

10. Mareedu, S., Pachon, R., Thilagavathi, J., Fefelova, N., Balakrishnan, R., Niranjan, N., Xie, L.H., and Babu, G.J. (2021). Sarcolipin haploinsufficiency prevents dystrophic cardiomyopathy in mdx mice. Am. J. Physiol. - Hear. Circ. Physiol. 320, H200–H210. 10.1152/AJPHEART.00601.2020.

11. Maurya, S.K., Bal, N.C., Sopariwala, D.H., Pant, M., Rowland, L.A., Shaikh, S.A., and Periasamy, M. (2015). Sarcolipin Is a Key Determinant of the Basal Metabolic Rate, and Its Overexpression Enhances Energy Expenditure and Resistance against Diet-induced Obesity. J. Biol. Chem. 290, 10840–10849. 10.1074/jbc.M115.636878.

12. Liu, Z., Zhang, Y., Qiu, C., Zhu, H., Pan, S., Jia, H., Kang, H., Guan, G., Hui, R., Zhu, L., et al. (2020). Diabetes mellitus exacerbates post-myocardial infarction heart failure by reducing sarcolipin promoter methylation. ESC Hear. Fail. 7, 1935–1948. 10.1002/ehf2.12789.

13. Briggs, F.N., Lee, K.F., Wechsler, A.W., and Jones, L.R. (1992). Phospholamban expressed in slow-twitch and chronically stimulated fast-twitch muscles minimally affects calcium affinity of sarcoplasmic reticulum Ca(2+)-ATPase. J. Biol. Chem. 267, 26056– 26061.

14. Rotter, D., Peiris, H., Grinsfelder, D.B., Martin, A.M., Burchfield, J., Parra, V., Hull, C., Morales, C.R., Jessup, C.F., Matusica, D., et al. (2018). Regulator of Calcineurin 1 helps coordinate whole-body metabolism and thermogenesis. EMBO Rep. 19, e44706. 10.15252/EMBR.201744706.

15. Habtemichael, E.N., Li, D.T., Paulo Camporez, J., Westergaard, X.O., Sales, C.I., Liu, X., López-Giráldez, F., DeVries, S.G., Li, H., Ruiz, D.M., et al. Insulin-stimulated endoproteolytic TUG cleavage links energy expenditure with glucose uptake. Nat. Metab. 10.1038/s42255-021-00359-x.

16. Yang, J., Goldstein, J.L., Hammer, R.E., Moon, Y.A., Brown, M.S., and Horton, J.D. (2001). Decreased lipid synthesis in livers of mice with disrupted Site-1 protease gene. Proc. Natl. Acad. Sci. U. S. A. 98, 13607–13612. 10.1073/pnas.201524598.

17. Kim, J.Y., Garcia-Carbonell, R., Yamachika, S., Zhao, P., Dhar, D., Loomba, R., Kaufman, R.J., Saltiel, A.R., and Karin, M. (2018). ER Stress Drives Lipogenesis and Steatohepatitis via Caspase-2 Activation of S1P. Cell 175, 133–145.e15. 10.1016/j.cell.2018.08.020.

18. Kondo, Y., Fu, J., Wang, H., Hoover, C., McDaniel, J.M., Steet, R., Patra, D., Song, J., Pollard, L., Cathey, S., et al. (2018). Site-1 protease deficiency causes human skeletal dysplasia due to defective inter-organelle protein trafficking. JCI Insight 3, e121596. 10.1172/JCI.INSIGHT.121596.

19. Patra, D., Xing, X., Davies, S., Bryan, J., Franz, C., Hunziker, E.B., and Sandell, L.J. (2007). Site-1 protease is essential for endochondral bone formation in mice. J. Cell Biol. 179, 687–700. 10.1083/jcb.200708092.

20. Mousa, M.G., Vuppaladhadiam, L., Kelly, M.O., Pietka, T., Ek, S., Shen, K.C., Meyer, G.A., Finck, B.N., and Brookheart, R.T. (2023). Site-1 protease inhibits mitochondrial respiration by controlling the TGF-β target gene Mss51. Cell Rep. 42. 10.1016/j.celrep.2023.112336.

21. Schweitzer, G.G., Gan, C., Bucelli, R.C., Wegner, D., Schmidt, R.E., Shinawi, M., Finck, B.N., and Brookheart, R.T. (2019). A mutation in Site-1 Protease is associated with a complex phenotype that includes episodic hyperCKemia and focal myoedema. Mol. Genet. Genomic Med. 7. 10.1002/mgg3.733.

22. Babu, G.J., Bhupathy, P., Carnes, C.A., Billman, G.E., and Periasamy, M. (2007). Differential expression of sarcolipin protein during muscle development and cardiac pathophysiology. J. Mol. Cell. Cardiol. 43, 215–222. 10.1016/j.yjmcc.2007.05.009.

23. Pereyra, A.S., Fernandez, R.F., Amorese, A., Castro, J.N., Lin, C.-T., Spangenburg, E.E., and Ellis, J.M. (2024). Loss of mitochondria long-chain fatty acid oxidation impairs skeletal muscle contractility by disrupting myofibril structure and calcium homeostasis. Mol. Metab. 89, 102015. 10.1016/j.molmet.2024.102015.

24. Tupling, A.R., Asahi, M., and MacLennan, D.H. (2002). Sarcolipin Overexpression in Rat Slow Twitch Muscle Inhibits Sarcoplasmic Reticulum Ca2+ Uptake and Impairs Contractile Function. J. Biol. Chem. 277, 44740–44746. 10.1074/jbc.M206171200.

25. Fajardo, V.A., Gamu, D., Mitchell, A., Bloemberg, D., Bombardier, E., Chambers, P.J., Bellissimo, C., Quadrilatero, J., and Tupling, A.R. (2017). Sarcolipin deletion exacerbates soleus muscle atrophy and weakness in phospholamban overexpressing mice. PLoS One 12, e0173708. 10.1371/journal.pone.0173708.

26. Ottenheijm, C.A.C., Hidalgo, C., Rost, K., Gotthardt, M., and Granzier, H. (2009). Altered contractility of skeletal muscle in mice deficient in titin’s M-band region. J. Mol. Biol. 393, 10–26. 10.1016/j.jmb.2009.08.009.

27. Burkholder, T.J., Fingado, B., Baron, S., and Lieber, R.L. (1994). Relationship between muscle fiber types and sizes and muscle architectural properties in the mouse hindlimb. J. Morphol. 221, 177–190. 10.1002/jmor.1052210207.

28. Tupling, A.R., Bombardier, E., Gupta, S.C., Hussain, D., Vigna, C., Bloemberg, D., Quadrilatero, J., Trivieri, M.G., Babu, G.J., Backx, P.H., et al. (2011). Enhanced ca 2 transport and muscle relaxation in skeletal muscle from sarcolipin-null mice. Am. J. Physiol. - Cell Physiol. 301, 841–849. 10.1152/ajpcell.00409.2010.

29. Ye, J. (2020). Transcription factors activated through RIP (regulated intramembrane proteolysis) and RAT (regulated alternative translocation). J. Biol. Chem. 295, 10271– 10280. 10.1074/jbc.REV120.012669.

30. Zeng, L., Lu, M., Mori, K., Luo, S., Lee, A.S., Zhu, Y., and Shyy, J.Y.J. (2004). ATF6 modulates SREBP2-mediated lipogenesis. EMBO J. 23, 950–958. 10.1038/SJ.EMBOJ.7600106/ASSET/462ADAC0-A57D-4DFF-9E0E-DD76FC590A02/ASSETS/GRAPHIC/EMBJ7600106-FIG-0008-M.JPG.

31. Wang, Y., Vera, L., Fischer, W.H., and Montminy, M. (2009). The CREB coactivator CRTC2 links hepatic ER stress and fasting gluconeogenesis. Nature 460, 534–537. 10.1038/nature08111.

32. Blackwood, E.A., Hofmann, C., Santo Domingo, M., Bilal, A.S., Sarakki, A., Stauffer, W., Arrieta, A., Thuerauf, D.J., Kolkhorst, F.W., Müller, O.J., et al. (2019). ATF6 Regulates Cardiac Hypertrophy by Transcriptional Induction of the mTORC1 Activator, Rheb. Circ. Res. 124, 79–93. 10.1161/CIRCRESAHA.118.313854.

33. Wu, J., Rutkowski, D.T., Dubois, M., Swathirajan, J., Saunders, T., Wang, J., Song, B., Yau, G.D.-Y., and Kaufman, R.J. (2007). ATF6alpha optimizes long-term endoplasmic reticulum function to protect cells from chronic stress. Dev. Cell 13, 351–364. 10.1016/j.devcel.2007.07.005.

34. Ye, J., Rawson, R.B., Komuro, R., Chen, X., Davé, U.P., Prywes, R., Brown, M.S., and Goldstein, J.L. (2000). ER stress induces cleavage of membrane-bound ATF6 by the same proteases that process SREBPs. Mol. Cell 6, 1355–1364. 10.1016/S1097-2765(00)00133-7.

35. Belmont, P.J., Tadimalla, A., Chen, W.J., Martindale, J.J., Thuerauf, D.J., Marcinko, M., Gude, N., Sussman, M.A., and Glembotski, C.C. (2008). Coordination of growth and endoplasmic reticulum stress signaling by regulator of calcineurin 1 (RCAN1), a novel ATF6-inducible gene. J. Biol. Chem. 283, 14012–14021. 10.1074/jbc.M709776200.

36. Chen, X., Shen, J., and Prywes, R. (2002). The luminal domain of ATF6 senses endoplasmic reticulum (ER) stress and causes translocation of ATF6 from the er to the Golgi. J. Biol. Chem. 277, 13045–13052. 10.1074/jbc.M110636200.

37. Du, K., Asahara, H., Jhala, U.S., Wagner, B.L., and Montminy, M. (2000). Characterization of a CREB Gain-of-Function Mutant with Constitutive Transcriptional Activity In Vivo. Mol. Cell. Biol. 20, 4320–4327. 10.1128/mcb.20.12.4320-4327.2000.

38. Gorski, P.A., Jang, S.P., Jeong, D., Lee, A., Lee, P., Oh, J.G., Chepurko, V., Yang, D.K., Kwak, T.H., Eom, S.H., et al. (2019). Role of SIRT1 in Modulating Acetylation of the Sarco-Endoplasmic Reticulum Ca2+-ATPase in Heart Failure. Circ. Res. 124, e63–e80. 10.1161/CIRCRESAHA.118.313865.

39. Kho, C., Lee, A., Jeong, D., Oh, J.G., Chaanine, A.H., Kizana, E., Park, W.J., and Hajjar, R.J. (2011). SUMO1-dependent modulation of SERCA2a in heart failure. Nature 477, 601–605. 10.1038/nature10407.

40. Gramolini, A.O., Trivieri, M.G., Oudit, G.Y., Kislinger, T., Li, W., Patel, M.M., Emili, A., Kranias, E.G., Backx, P.H., and Maclennan, D.H. (2006). Cardiac-specific overexpression of sarcolipin in phospholamban null mice impairs myocyte function that is restored by phosphorylation. Proc. Natl. Acad. Sci. U. S. A. 103, 2446–2451. 10.1073/pnas.0510883103.

41. Kaspari, R.R., Reyna-Neyra, A., Jung, L., Torres-Manzo, A.P., Hirabara, S.M., and Carrasco, N. (2020). The paradoxical lean phenotype of hypothyroid mice is marked by increased adaptive thermogenesis in the skeletal muscle. Proc. Natl. Acad. Sci. U. S. A. 117, 22544–22551. 10.1073/pnas.2008919117.

42. Myers, J.W., Park, W.Y., Eddie, A.M., Shinde, A.B., Prasad, P., Murphy, A.C., Leonard, M.Z., Pinette, J.A., Rampy, J.J., Montufar, C., et al. (2024). Systemic inhibition of de novo purine biosynthesis prevents weight gain and improves metabolic health by increasing thermogenesis and decreasing food intake. bioRxiv Prepr. Serv. Biol. 10.1101/2024.10.28.620705.

43. Picard, M., Hepple, R.T., and Burelle, Y. (2012). Mitochondrial functional specialization in glycolytic and oxidative muscle fibers: Tailoring the organelle for optimal function. Am. J. Physiol. - Cell Physiol. 302, 629–641. 10.1152/AJPCELL.00368.2011/ASSET/IMAGES/LARGE/ZH00021268360003.JPEG.

44. McMillan, E.M., and Quadrilatero, J. (2011). Differential apoptosis-related protein expression, mitochondrial properties, proteolytic enzyme activity, and DNA fragmentation between skeletal muscles. Am. J. Physiol. Regul. Integr. Comp. Physiol. 300, R531–43. 10.1152/ajpregu.00488.2010.

45. Picard, M., Csukly, K., Robillard, M.-E., Godin, R., Ascah, A., Bourcier-Lucas, C., and Burelle, Y. (2008). Resistance to Ca2+-induced opening of the permeability transition pore differs in mitochondria from glycolytic and oxidative muscles. Am. J. Physiol. Regul. Integr. Comp. Physiol. 295, R659–68. 10.1152/ajpregu.90357.2008.

46. Fujita, R., Yoshioka, K., Seko, D., Suematsu, T., Mitsuhashi, S., Senoo, N., Miura, S., Nishino, I., and Ono, Y. (2018). Zmynd17 controls muscle mitochondrial quality and whole-body metabolism. FASEB J. 32, 5012–5025. 10.1096/fj.201701264R.

47. Guillet-Deniau, I., Mieulet, V., Lay, S. Le, Achouri, Y., Carre, D., Girard, J., Foufelle, F., and Ferre, P. (2002). Sterol Regulatory Element Binding Protein-1c Expression and Action in Rat Muscles: Insulin-Like Effects on the Control of Glycolytic and Lipogenic Enzymes and UCP3 Gene Expression. Diabetes 51, 1722–1728. 10.2337/diabetes.51.6.1722.

48. Seo, H.Y., Kim, M.K., Min, A.K., Kim, H.S., Ryu, S.Y., Kim, N.K., Lee, K.M., Kim, H.J., Choi, H.S., Lee, K.U., et al. (2010). Endoplasmic reticulum stress-induced activation of activating transcription factor 6 decreases cAMP-stimulated hepatic gluconeogenesis via inhibition of CREB. Endocrinology 151, 561–568. 10.1210/en.2009-0641.

49. Patel, K.H., Talovic, M., Dunn, A.J., Patel, A., Vendrell, S., Schwartz, M., and Garg, K. (2020). Aligned nanofibers of decellularized muscle extracellular matrix for volumetric muscle loss. J. Biomed. Mater. Res. B. Appl. Biomater. 108, 2528–2537. 10.1002/jbm.b.34584.

50. Wang, Y., Shen, J., Arenzana, N., Tirasophon, W., Kaufman, R.J., and Prywes, R. (2000). Activation of ATF6 and an ATF6 DNA binding site by the endoplasmic reticulum stress response. J. Biol. Chem. 275, 27013–27020. 10.1074/jbc.M003322200.

51. Liu, Q., Busby, J.C., and Molkentin, J.D. (2009). Interaction between TAK1-TAB1-TAB2 and RCAN1-calcineurin defines a signalling nodal control point. Nat. Cell Biol. 11, 154–161. 10.1038/NCB1823.

52. Clipstone, N.A., Fiorentino, D.F., and Crabtree, G.R. (1994). Molecular analysis of the interaction of calcineurin with drug-immunophilin complexes. J. Biol. Chem. 269, 26431– 26437. 10.1016/S0021-9258(18)47212-2.

53. DeBose-Boyd, R.A., Brown, M.S., Li, W.P., Nohturfft, A., Goldstein, J.L., and Espenshade, P.J. (1999). Transport-dependent proteolysis of SREBP: relocation of site-1 protease from Golgi to ER obviates the need for SREBP transport to Golgi. Cell 99, 703– 712.

54. Geromella, M.S., Braun, J.L., and Fajardo, V.A. (2023). Measuring SERCA-mediated calcium uptake in mouse muscle homogenates. STAR Protoc. 4, 101987. 10.1016/j.xpro.2022.101987.

