## Supplemental Figure 1 for "Site-1 Protease is a negative regulator of sarcolipin promoter activity"

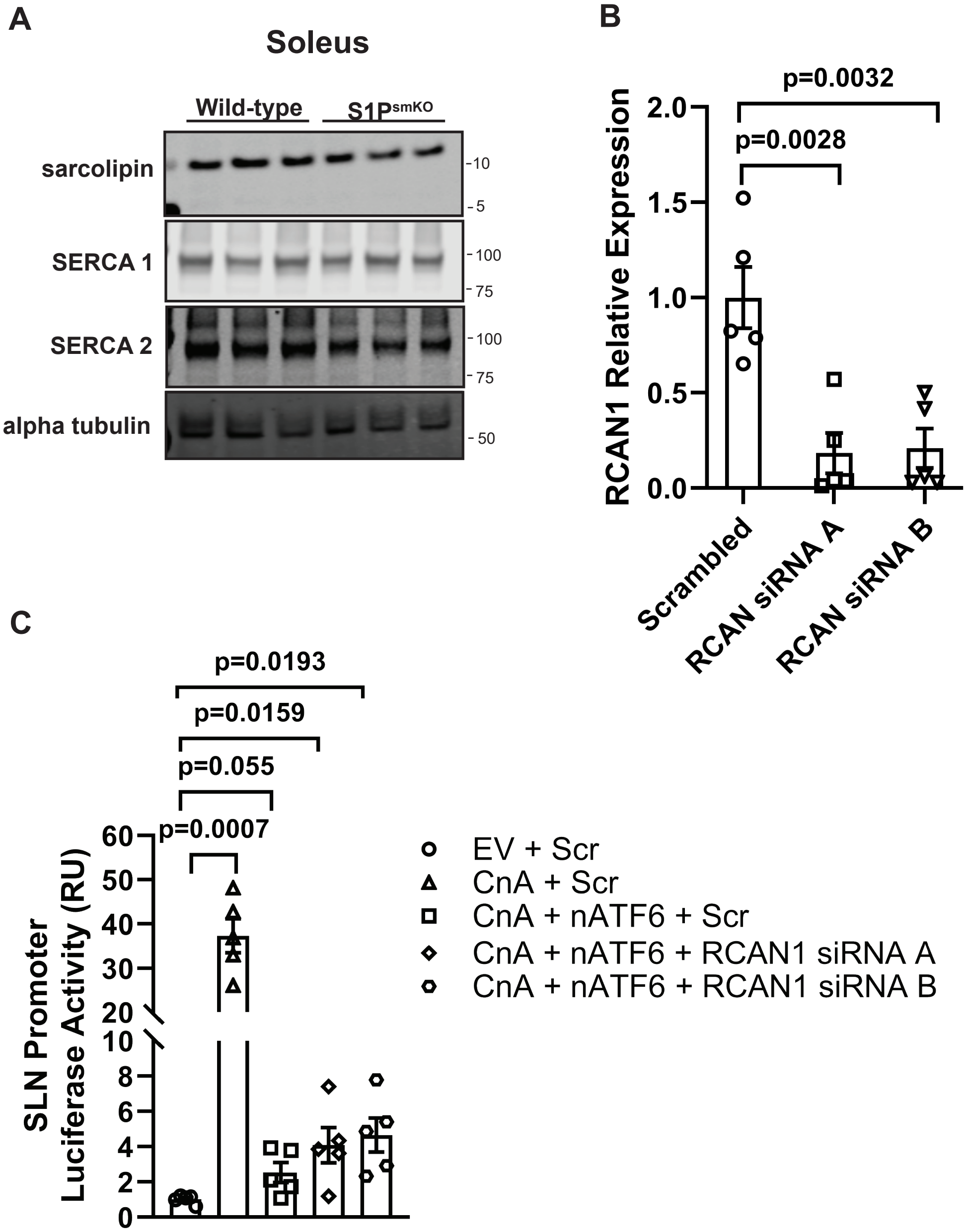


**Supplemental Figure 1.** (A) Western blot of sarcolipin protein levels in soleus muscles of wild-type and S1PsmKO mice. n=3/group. (B) RCAN1 mRNA levels in scrambled and RCAN1 siRNA transfected C2C12 cells. n=5/group. (C) Sarcolipin promoter activity in cells transfected with EV, CnA, or nATF6 plus scrambled or RCAN1 siRNAs as indicated. n=5/group. EV, empty vector; CnCA, constitutively active calcineurin, nATF6, active ATF6. Data are reported as ± SEM.
