## Supplemental Table 1 for "Site-1 Protease is a negative regulator of sarcolipin promoter activity"

Primer Sequences

| Gene | Forward (5'-3') | Reverse (5'-3') |
|---|---|---|
| Sarcolipin | tcc tcg tga ggt cct acc aa | tag agc att gga agc tcg gg |
| 36B4 | gca gac aac gtg ggc tcc aag cag at | ggt cct cct tgg tga aca cga agc cc |
| RCAN1 | gat gga gga ggt gga tct gc | ttc aaa ttt ggc ccg gca c |
